# A Novel Surgical Method for Continuous Intra-Portal Infusion of Gut Microbial Metabolites in Mice

**DOI:** 10.1101/2020.10.29.360628

**Authors:** Danny Orabi, Lucas J. Osborn, Kevin Fung, Federico Aucejo, Ibrahim Choucair, Beckey DeLucia, Zeneng Wang, Jan Claesen, J. Mark Brown

**Author notes:** Contributed equally to this work and are to be considered co-first authors (Danny Orabi, Lucas J. Osborn). To whom correspondence should be addressed: Department of Cardiovascular and Metabolic Sciences, Cleveland Clinic, Cleveland, OH 44195, USA.

## Abstract

Gut microbial-derived metabolites have been shown to play key roles in human physiology and disease. However, establishing mechanistic links between gut microbial metabolites and disease pathogenesis in animal models presents many challenges. The major route of absorption for microbe-derived small molecules is venous drainage via the portal vein to the liver. In the event of extensive liver first pass- or presystemic hepatic metabolism, the route of administration of these metabolites becomes critical. Here we describe a novel portal vein cannulation technique using a subcutaneously implanted osmotic pump to achieve continuous portal vein infusion in mice. First, the microbial metabolite trimethylamine (TMA) was administered over 4 weeks and compared to a vehicle control. Using a liquid chromatography-tandem mass spectrometry (LC-MS/MS), an increase in peripheral plasma levels of TMA and its host liver-derived co-metabolite trimethylamine-N-oxide (TMAO) were observed in a sexually-dimorphic manner. Next, 4-hydroxyphenylacetic acid (4-HPAA), a structurally distinct microbial metabolite that undergoes extensive hepatic first pass metabolism, was administered intraportally to examine effects on hepatic gene expression. As expected, there was no difference in peripheral plasma 4-HPAA levels yet liver tissue demonstrated higher levels of 4-HPAA when compared to the control group. More importantly, significant changes were observed in hepatic gene expression using an unbiased RNA sequencing approach. Collectively, this work describes a novel method for administering gut microbe-derived metabolites via the portal vein, mimicking their physiologic delivery *in vivo*.

**Importance:** Recent efforts have underscored the importance of the gut microbial community as a meta-endocrine organ impacting host physiology through systemic delivery of gut-microbial metabolites [Brown and Hazen, 2015]. Microbial metabolites are first delivered to the liver via the portal vein following venous drainage of the gastrointestinal tract. This route of absorption is often crucial by allowing the liver to biotransfrom these molecules prior to entering the peripheral circulation. Microbial metabolites are frequently studied in animal models by incorporation into diet or drinking water. This method falls short as inconsistent oral intake, inconsistent gastrointestinal absorption, and further modification of the metabolite by gut microbes yield imprecise levels of drug delivery. In efforts to overcome this, the physiological impact of microbial metabolites is often studied by intermittent exogenous administration of a metabolite in a non-physiologically relevant manner such as intravenous injection, intraperitoneal injection, or subcutaneous administration, all placing a relatively large proportion of the metabolite directly into the peripheral circulation. Although these approaches can effectively raise circulating metabolites levels in some cases, they do not mimic the natural delivery of gut microbial-derived small molecules through the portal circulation to the liver. Here we describe a novel surgical method to continuously deliver precise amounts of gut microbial metabolites intraportally to better recapitulate the natural systemic delivery route of microbial metabolites to the liver. This model will improve the interrogation of gut microbial metabolites and their associations to disease by providing an unmatched level of resolution when manipulating the portal blood metabolome.

## Introduction

Gut microbes are key modulators of human physiology and are well appreciated as primary producers of metabolites that have been shown to impact human health [Glowacki and Martens, 2020]. Disease states including cardiovascular disease (CVD), chronic kidney disease (CKD), diabetes, cancer, and cancer therapeutic efficacy, all contain a demonstrable microbial metabolite component [Wang et al., 2011, Gupta et al., 2020, Koh et al., 2018, Nougayrède et al., 2006, Mager et al., 2020]. However, most mechanisms by which gut microbial metabolites impact the host are poorly understood. Furthermore, a limiting factor in the mechanistic interrogation of microbial metabolites is the lack of physiologically relevant delivery mechanisms.

The canonical route of absorption for molecules originating from the gastrointestinal tract is venous drainage via the portal vein to the liver. This route of absorption is often crucial by allowing the liver to biotransform these molecules prior to entering the peripheral circulation. In many cases, molecules delivered via the portal vein to the liver are subjected to extensive presystemic hepatic metabolism, also known as the liver first pass effect, such that their biotransformed products are the predominant forms seen in the peripheral circulation [Pond and Tozer, 1984]. It is well appreciated that the portal blood metabolome can impact hepatic physiology in the context of numerous diseases such as alcoholic liver disease (ALD), non-alcoholic fatty liver disease (NAFLD), and hepatocellular carcinoma (HCC) [Lang et al., 2020, Yoshimoto et al., 2013]. As such, the administration route of gut-derived metabolites becomes critical, where delivery of any particular metabolite should be done in a manner that recapitulates the natural *in vivo* delivery. Currently, common routes of administration include intravenous, intraperitoneal, and subcutaneous injections, through intermittent oral gavage, or by incorporation into the diet or drinking water. Critically, none of these approaches ensure that the metabolite of interest remains unmodified by intestinal or gut microbial metabolism as it passes through portal circulation to the liver, in order to mimic normal physiologic conditions.

The dietary nutrients choline, phosphatidylcholine, and carnitine can serve as precursors for bacterial trimethylamine (TMA) generation which is subject to further hepatic oxidation by flavin monooxygenases [Koeth et al., 2013, Wang et al., 2011]. This two-step meta-organismal metabolic process yields the pro-inflammatory, proatherosclerotic, and prothrombotic metabolite trimethylamine-N-oxide (TMAO) [Koeth et al., 2013, Tang et al., 2013, Wang et al., 2011, Zhu et al., 2016]. Due to the well-studied nature of the TMA/TMAO pathway and the robust characterization of murine plasma TMA and TMAO pools, we selected TMA as a candidate molecule for a proof of concept study for our novel surgical procedure for continuous intraportal infusion. Additionally, TMA has a relatively low rate of first-pass hepatic metabolism which is evident from the ability to manipulate peripheral plasma TMA through incorporation of its dietary precursors into rodent chow [Koeth et al., 2013]. Consequently, TMA presented itself as an ideal candidate molecule for a preliminary proof of concept study given its ability to be detected in peripheral plasma resulting from low first-pass hepatic metabolism.

Gut microbial metabolism of dietary input yields a portal blood metabolome rich in microbe-derived metabolites that are delivered to the liver via portal circulation [Marin et al., 2015, Lang and Schnabl, 2020]. Polyphenols are abundant in fruits and vegetables and are known to undergo extensive microbial metabolism [Marin et al., 2015, Braune and Blaut, 2016]. Although polyphenolic flavonoids alone have demonstrated efficacy in the context of various inflammatory disorders [Assini et al., 2013, Burke et al., 2018, Liu et al., 2017], flavonoid-mediated attenuation of disease in disorders such as arthritis, obesity, and atherosclerosis has been reported to be microbe-dependent [Aa et al., 2019, Esposito et al., 2015, Wang et al., 2010]. The dietary flavonol kaempferol, has been described to undergo specific bacterial cleavage of its C-ring to yield a microbial flavonoid metabolite, 4-hydroxyphenylacetic acid (4-HPAA) [Griffiths and Smith, 1972, Winter et al., 1991]. Of clinical relevance, Hoyles and colleagues reported 4-HPAA as being negatively correlated with free fat mass, waist circumference, and area under the curve for an oral glucose tolerance test in a study of 105 obese Caucasian females [Hoyles et al., 2018]. A key characteristic of 4-HPAA that warranted its inclusion in this study, is that it undergoes rapid elimination following intravenous injection [Zabela et al., 2016]. In order to study a metabolite with such a short half-life, continuous delivery is required to more fully recapitulate *in vivo* conditions.

Herein, we describe a novel surgical approach to continuously administer microbial and other gut-derived metabolites intraportally. We hypothesized that this technique would allow interrogation of the impact that microbial metabolites have on host physiology. Our findings suggest that the continuous intraportal infusion of TMA is sufficient to significantly raise plasma levels of both TMA and its primary hepatic metabolite TMAO in mice when compared to single-housed, sham, and saline control groups. Moreover, we demonstrate that continuous infusion of a metabolite with high first-pass metabolism, 4-HPAA, accumulates in the liver while both the free and total plasma concentrations remain unchanged. Additionally, the continuous infusion of 4-HPAA is sufficient to alter the hepatic transcriptional landscape when compared to the vehicle control group.

## Results

### A novel surgical procedure for the continuous intraportal infusion of microbial metabolites

The surgical procedure in its entirety is stepwise illustrated in Figure 1, and is described in detail in the methods section. It is critical to ensure that strict adherence to sterile technique is maintained for the duration of the procedure given this is a survival surgery designed for long-term metabolite infusion. First, a laparotomy is performed, the intestines are externalized, the duodenum is rotated (Figure 1B), and the left liver lobe is externalized to expose the portal vein (Figure 1C) (Supplementary video 1). Next, a polyurethane catheter is advanced within the portal vein using a Teflon-coated guidewire to facilitate entry and is secured to the portal vein with a microsuture (Figures 1D – 1I) (Supplementary video 2). The catheter is then externalized from the peritoneal cavity, where it is connected to a mini osmotic pump (Figure 1J) (Supplementary video 3). The osmotic pump is placed within the subcutaneous tissue of the interscapular space on the dorsum of the mouse and the abdominal wall and skin are closed (Figures 1K, 1L). One week after surgery, ultrasound imaging demonstrated the tip of the cannula properly positioned within the portal vein and surrounding liver-directed laminar blood flow (Figure (Supplementary videos 4, 5). At the completion of the experiment, mice were placed under isoflurane anesthesia and saline was gently injected through the cannula to ensure catheter patency and position (Supplementary video 6).

**Figure 1.**
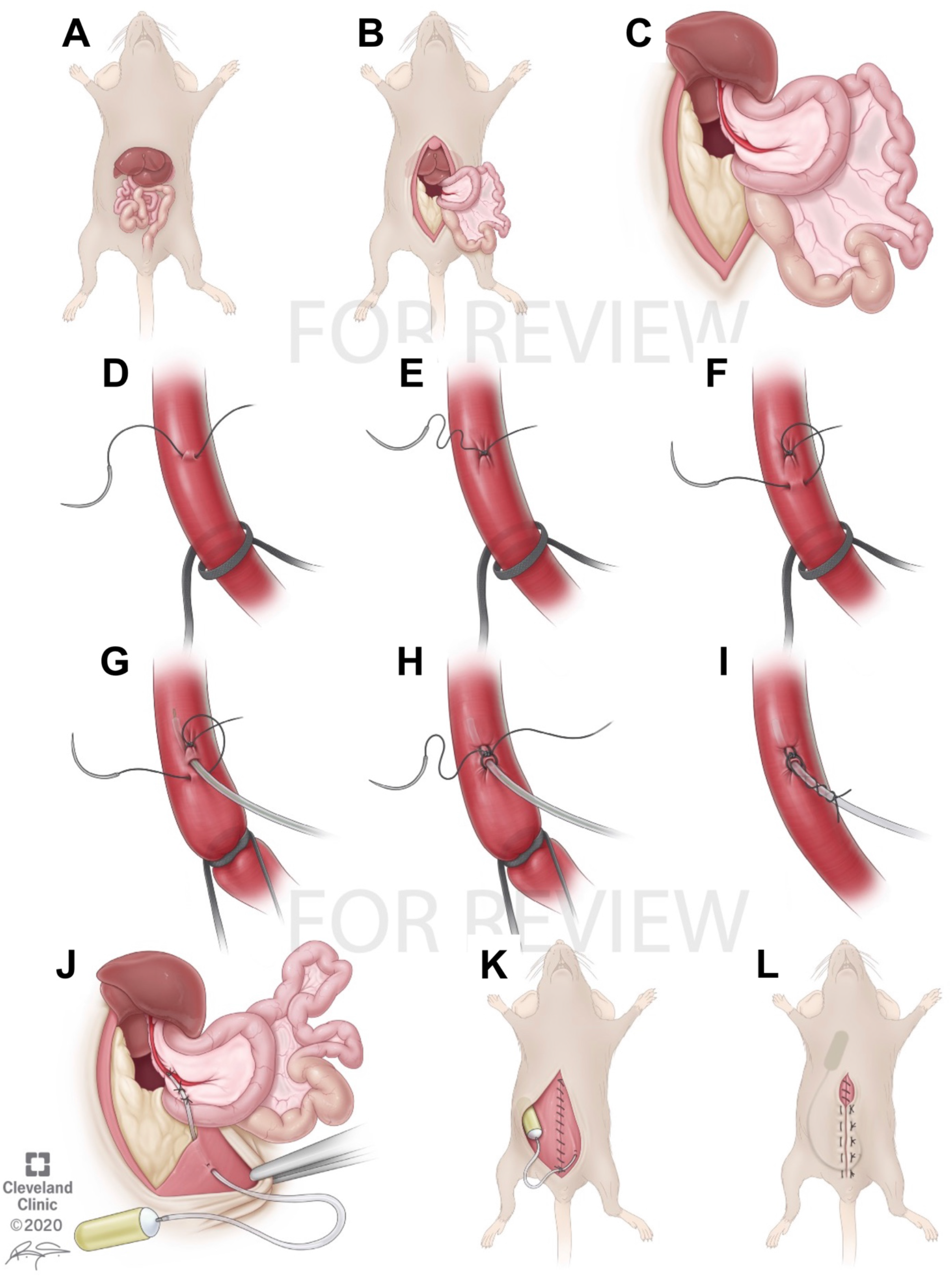
Stepwise illustration of portal vein cannulation procedure. (A) Abdominal anatomy. (B) Midline incision and laparotomy, with leftward externalization of intestines and overlying duodenum. (C) Externalization of left liver lobe superiorly. (D-F) Proximal vessel loop is loose. Using a 10-0 nylon micro suture, a stitch is placed in the anterior one-fourth of the vein 3 mm distal to the vessel loop. The needle from this stitch is placed in the anterior one-fourth of the vein 1 mm proximal to the stitch. (G) The vessel loop is pulled taught. The guidewire is used to facilitate insertion. (H) The guidewire is withdrawn. The suture ends are tied together, slightly crimping the catheter. (I) The vessel loop is loosened and cut. Each of the tails of the suture are wrapped around the catheter twice and tied to create a “Chinese finger trap” effect. (J) Two anchor stitches are placed in the mesentery of the duodenum to secure the catheter. The catheter is externalized from the left lower quadrant and anchored to the peritoneal side of the abdominal wall. (K) The abdominal wall is closed, and the catheter anchored to the inferior aspect of the closure. (L) The osmotic pump is placed in the interscapular subcutaneous space. The skin is closed.

**Figure 2.**
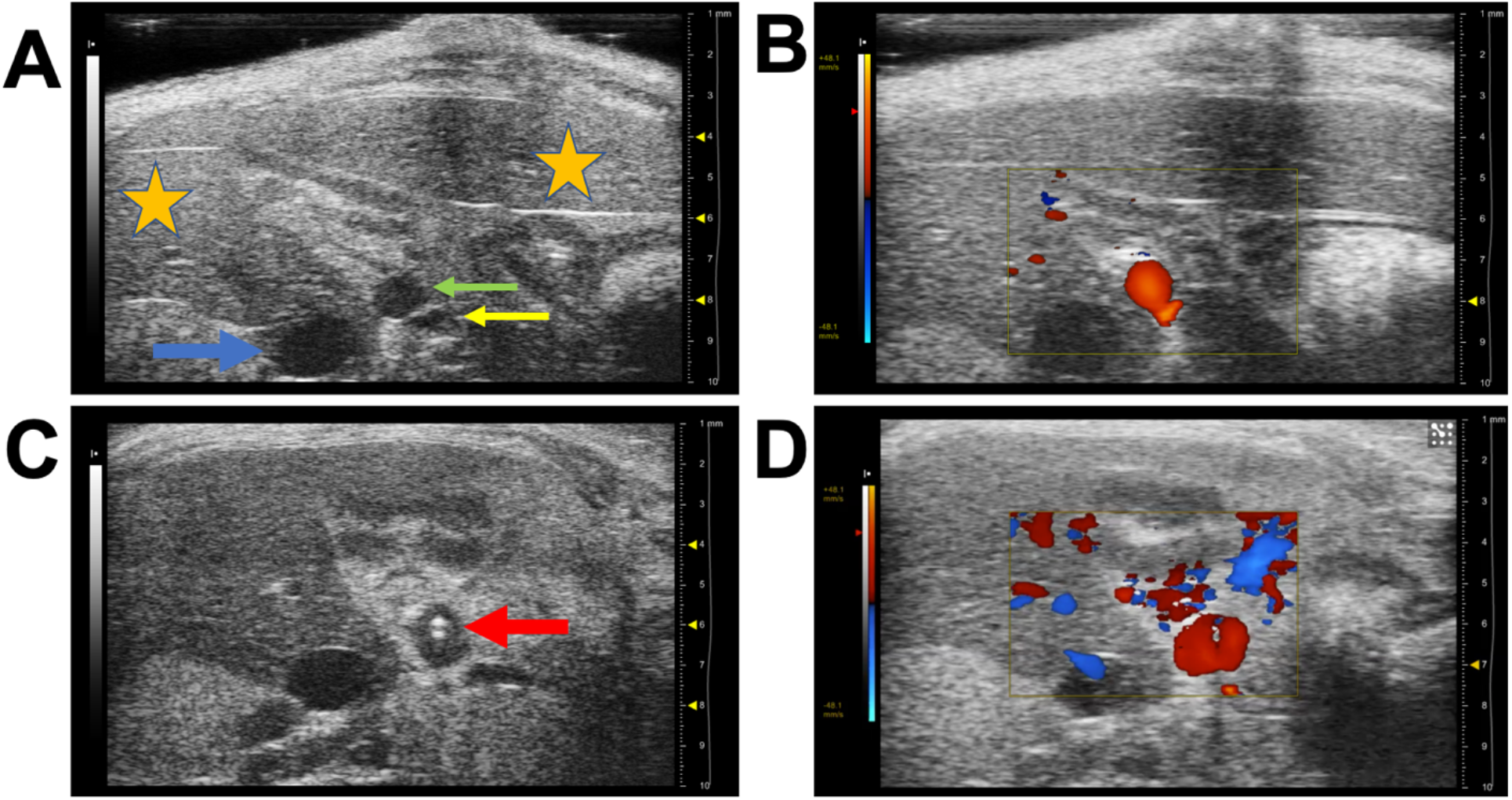
Ultrasound imaging of catheterized portal vein. Mice underwent ultrasound imaging one week after surgery (n=3). (A) The portal vein proximal to the catheter appeared patent and non-dilated, and (B) demonstrated liver-directed laminar flow on doppler imaging. (C) The tip of the catheter was visualized within the distal portal vein (red arrow), and (D) liver-directed laminar flow was visualized around the catheter. [Orange star: liver; green arrow: portal vein; yellow arrow: hepatic artery; blue arrow: inferior vena cava.]

### Continuous infusion of TMA provides proof of concept for the surgical procedure

As a preliminary proof of concept, we administered TMA continuously for 4 weeks in both male (Figure 3A-C) and female (Figure 3D-F) C57BL/6 mice. As the post-surgery recovery requires each mouse to be single-housed, we included a single housed control group (SH) when measuring plasma TMA and TMAO. Additionally, we included sham surgery control animals that were opened via midline laparotomy and had their portal vein dissected but not cannulated (Sham). We also included control animals that had their portal vein cannulated and received normal saline from the osmotic pump for the duration of the study (NS). As expected, the continuous intraportal infusion of TMA over 4 weeks led to a significant increase in peripheral plasma TMA levels in male mice when compared to saline control animals as measured by LC-MS/MS (Figure 3A, 3C). Peripheral plasma TMAO was not significantly different between the male mice receiving TMA or saline (Figure 3B, 3C). Conversely, female mice had appreciably higher peripheral plasma concentrations of TMA when compared to the normal saline control (Figure 3D, 3F), as well as markedly higher levels of the host liver-derived co-metabolite TMAO (Figure 3E, 3F). These data are further corroborated by previous studies demonstrating that female mice have >1,000-fold higher expression of FMO3, the main enzyme in the liver responsible for the conversion of TMA to TMAO [Wang et al., 2011, PMID: 23312283]. Hence, the higher TMA:TMAO ratio in males and concomitantly lower TMA:TMAO ratio in females was expected. Collectively, these data provide proof of concept that the surgical model of continuous intraportal infusion of microbial metabolites is feasible.

**Figure 3.**
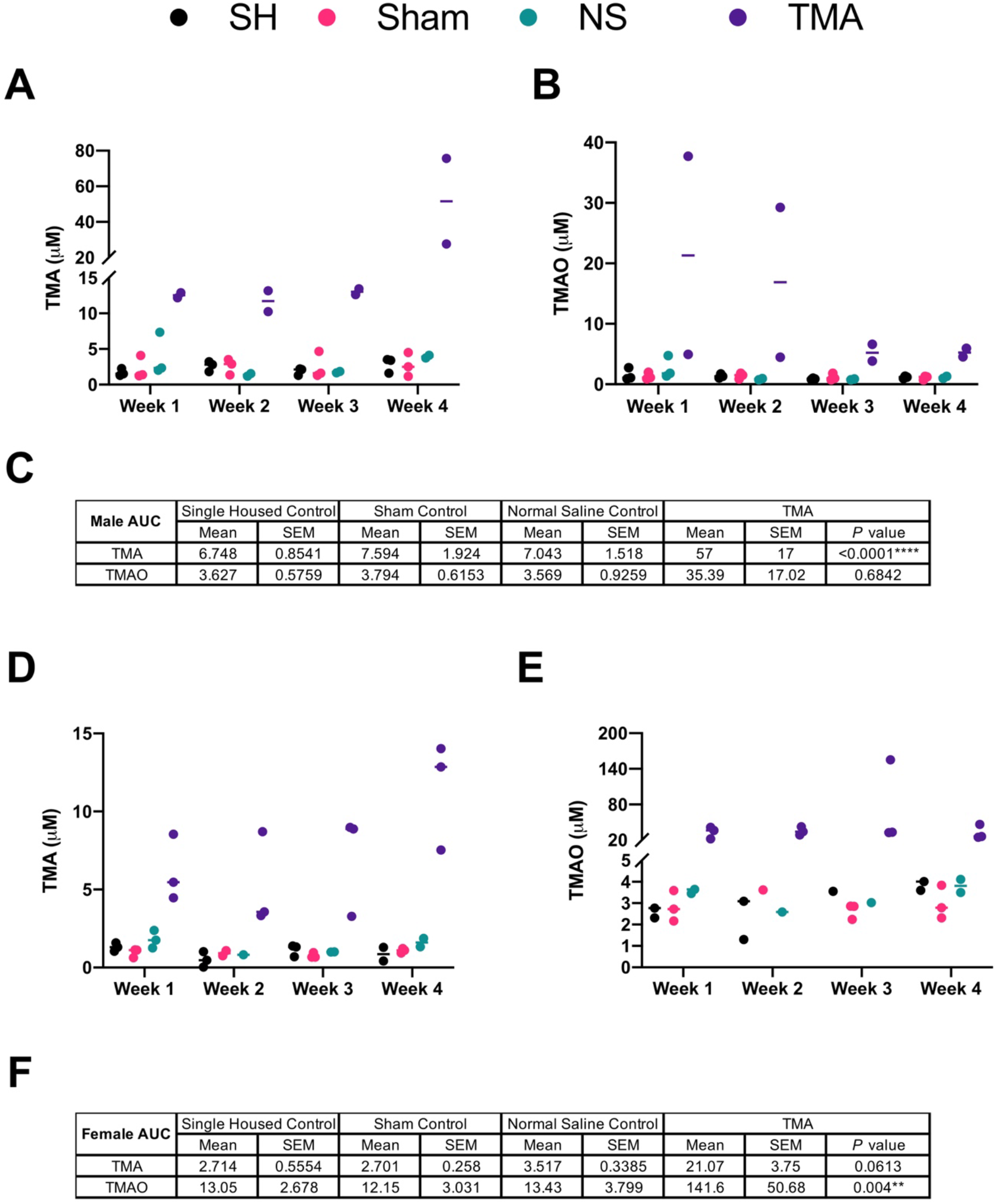
Continuous infusion of TMA– a microbial metabolite with low hepatic first-pass metabolism. (A) Peripheral plasma levels of trimethylamine (TMA) in male C57BL/6 mice receiving TMA via intraportal catheter (n=2-3). Shown also are single-housed controls (SH), sham surgery controls (Sham), and osmotic pump normal saline controls (NS). (B) Peripheral plasma trimethylamine-N-oxide (TMAO) concentration in male C57BL/6 mice receiving TMA or normal saline via intraportal catheter with relevant single-housed and sham controls. (C) Summary of area under the curve (AUC) mean and standard error of the mean (SEM) for the duration of the 4-week study. (D) Peripheral plasma levels of TMA in female C57BL/6 mice receiving TMA via intraportal catheter (n=2-3). Shown also are single-housed controls (SH), sham surgery controls (Sham), and osmotic pump normal saline controls (NS). (E) Peripheral plasma TMAO concentration in female C57BL/6 mice receiving TMA or normal saline via intraportal catheter with relevant single-housed and sham controls. (F) Summary of area under the curve (AUC) mean and standard error of the mean (SEM) for the duration of the 4-week study. *P* values were calculated using two-way ANOVA of saline controls compared to the TMA group with Dunnet’s correction for multiple comparisons.

### Continuous infusion of 4-HPAA– a microbe-derived metabolite with high first-pass hepatic metabolism

Next, we sought out to determine the efficacy of our surgical model using a metabolite with a rapid rate of hepatic clearance, 4-HPAA. Osmotic pumps with either normal saline or 4-HPAA dissolved in normal saline (Figure 4A) were implanted following the surgical procedure outlined above and were allowed to continuously infuse their contents for two weeks (Figure 4B). As expected due to the previously reported rapid clearance of 4-HPAA [Zabela et al., 2016], we did not observe any difference in either free or glucuronidated/sulfated levels of 4-HPAA in peripheral plasma between normal saline and 4-HPAA groups (Figure 4C). Moreover, we gained insight into the overall circulating pool of 4-HPAA where it was observed that 4-HPAA in plasma is either glucuronidated or sulfated (Glu/Sul) as measured by LC-MS/MS (Figure 4C). The liver is uniquely positioned to act as a central player in metabolite-mediated gut-liver crosstalk and ultimately controls the fate of many gut-derived metabolites. In support of this concept, a significant accumulation of free and glucuronidated/sulfated 4-HPAA was observed in the livers of mice receiving 4-HPAA compared to control mice (Figure 4D). In summary, these data suggest that much of the circulating 4-HPAA pool is glucuronidated/sulfated, plasma levels of 4-HPAA cannot be raised due to rapid hepatic clearance, and that accumulation of 4-HPAA occurs in the liver after continuous intraportal infusion.

**Figure 4.**
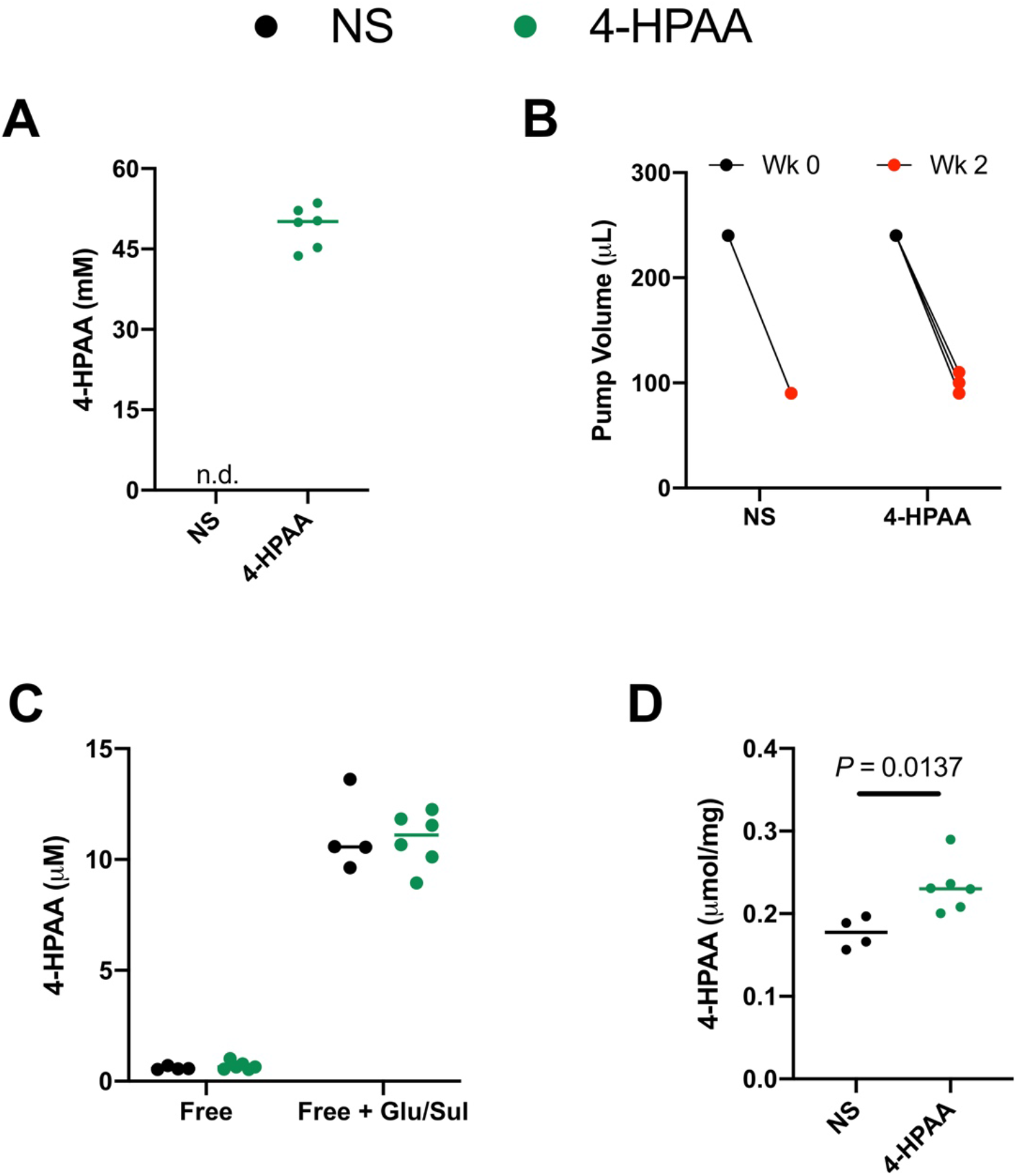
Continuous infusion of 4-HPAA– a microbial metabolite with high first-pass metabolism. (A) Osmotic pumps were filled with either normal saline (NS) or 4-hydroxyphenylacetic acid (4-HPAA). (B) Volume of either normal saline or 4-HPAA dispensed from the osmotic pumps after two weeks of continuous intraportal infusion (n=4-6). (C) A larger proportion of the 4-HPAA pool circulates in a glucuronidated/sulfated (Glu/Sul) form, reaching 10-fold higher total peripheral plasma 4-HPAA concentrations compared to the free 4-HPAA pool. (D) Significantly more 4-HPAA is detected in the liver (free + Glu/Sul), the last anatomic location where differential abundance of 4-HPAA is observed resulting from high first-pass hepatic metabolism of 4-HPAA. *P* values were calculated using an unpaired, two-tailed t-test.

### Physiologic administration of 4-HPAA into the portal vein alters the transcriptional landscape of the liver

Since 4-HPAA administration resulted in increased levels in the liver, we studied the compound’s local effects. We performed unbiased RNA sequencing in liver tissues from mice receiving continuous infusion of either saline or 4-HPAA (Figure 5). Non-metric multidimensional scaling (NMDS) and hierarchical clustering showed clear separation and predictable assignment of both saline and 4-HPAA groups (Figure 5A, 5B). The continuous infusion of this single microbial metabolite caused marked global hepatic transcriptional changes when compared to the normal saline control (Figure 5C). 4-HPAA caused the differential expression of several genes involved in the binding and uptake of ligands by scavenger receptors including alpha 1 microglobulin/bikunin precursor (*Ambp),* secreted acidic cysteine rich glycoprotein (*Sparc),* collectin sub-family member 11 (*Colec11),* apolipoprotein 7b (*Apol7b),* and hemoglobin, beta adult t chain (*Hbb-bt*) (Figure 5A, 5D). These unbiased RNA sequencing data suggest that the administration of a single microbial metabolite, 4-HPAA, is sufficient to alter the hepatic transcriptional landscape, providing the first clues into how the gut microbe-derived metabolite 4-HPAA can impact host liver gene expression.

**Figure 5.**
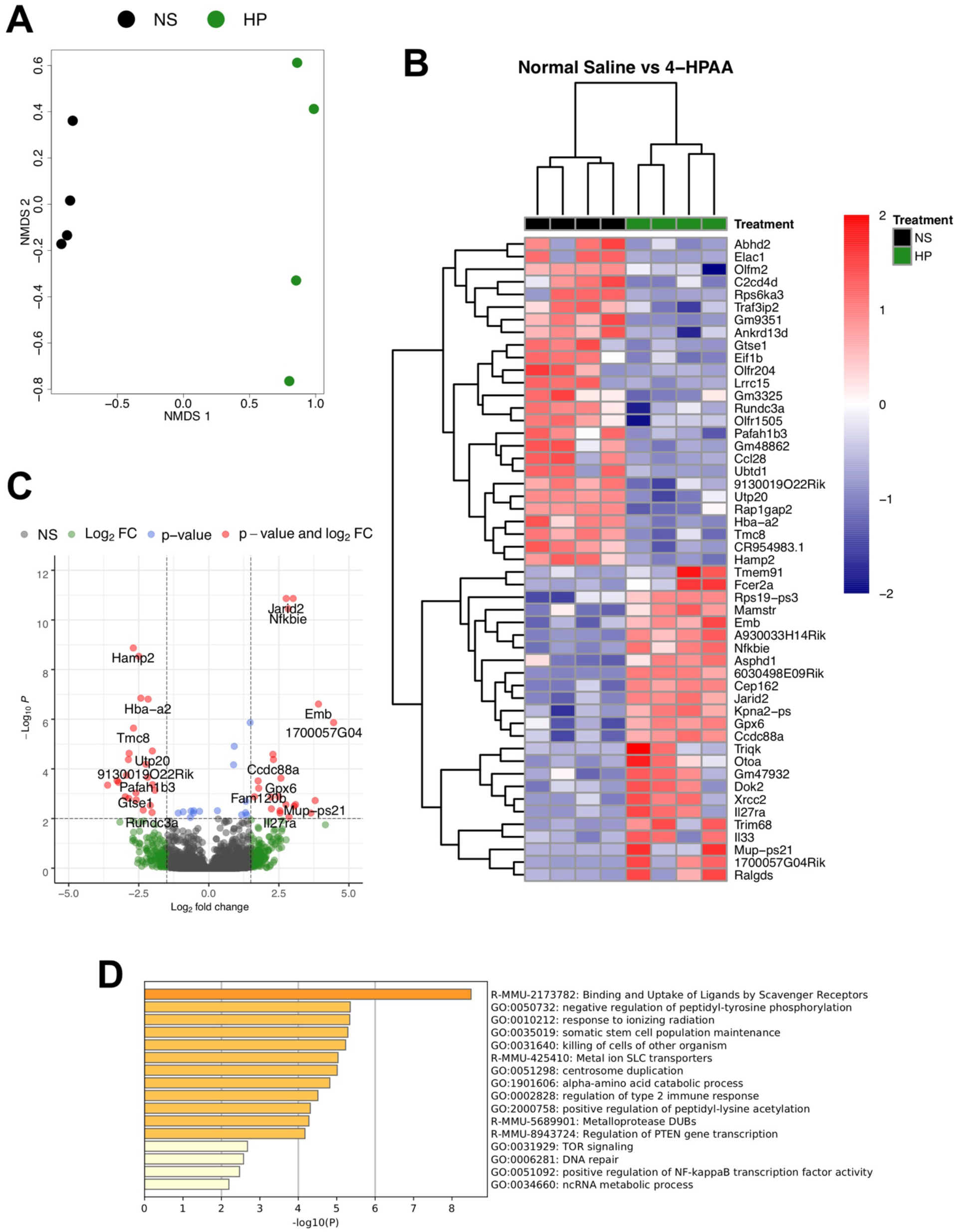
Continuous Portal infusion of 4-HPAA Changes the Hepatic Transcriptional Landscape. (A) Non-Metric Multi-Dimensional Scaling (NMDS) of RNA-Seq transcriptome data representing the hepatic gene expression signature of the top 500 differentially expressed transcripts between 4-HPAA treated mice (green) relative to normal saline control mice (black) as sorted by log2 fold change. The NMDS was performed using DESeq2 normalized counts. (B) Heatmap of hierarchically clustered differentially expressed genes arranged by adjusted *P* value and log2 fold change. Z-score normalized values scaled by row. (C) Volcano plot of RNA-Seq transcriptome data representing hepatic gene expression signature of 4-HPAA treated mice relative to normal saline control mice. Genes highlighted in red correspond to those that are significantly differentially expressed (Adjusted *P<*0.001) with a log2 fold change >1.5. (D) Gene ontology assignments of the top 250 differentially expressed genes as sorted by adjusted *P* value. N=4 per group for all RNASeq analysis.

## Discussion

Attempts to achieve long-term access to the portal circulation for continuous infusion or interval bolus administration have received attention over the last few decades. While mice are the most commonly used vertebrate in research, there are few examples of long-term administration of compounds into the portal circulation given the small size and fragility of the mouse portal vein. Rather, most portal vein cannulations are done in larger rat and porcine models [Strubbe et al. 1999; Oldhafer et al. 2020]. Our surgical technique is the first to use an indwelling pump along with portal vein cannulation to achieve continuous infusion over extended time periods. To our knowledge, the only two papers to describe portal vein continuous infusion in mice use an external swivel wheel and syringe pump with 5-foot long catheter tubing and infusion rate of 6 mL/day (1000-fold higher than our model) [Patijn et al 1998a, 1998b]. That model along with all others describing portal vein cannulation for interval bolus administration do not perform a confirmation procedure at necropsy to ensure proper intraportal position of the catheter tip [Shen et al. 2008; Vrancken Peeters et al 1996; Valentino et al 2013]. This is crucial as slight pressure on the sidewall of the portal vein will lead to erosion of the catheter into the vein sidewall or adjacent tissues. Additionally, all other models employ the intraperitoneal use of surgical glue at the catheter – portal vein insertion side, though glue is often non-sterile, produces an inflammatory reaction with dense adhesion formation, and affects gut peristalsis if accidentally applied on bowel [Bot et al 2010]. The method described here of portal vein cannulation and continuous infusion is novel in its ability to achieve precise administration of microbial metabolites in a physiologically relevant way. However, achieving long-term patency and the proper intraportal position of the catheter is technically challenging and it is the authors’ hope that the detail provided within will allow its more widespread use. The extensive subcutaneous dissection, length of surgery (approximately 50 minutes), and need to single-house animals should increase the thoughtfulness paid to a metabolite’s suitability for this technique (rapid hepatic first pass metabolism, involvement in enterohepatic circulation, etc.).

The method described herein provides the unique ability to study microbial metabolite-mediated gut-liver crosstalk with an unparalleled level of resolution in a physiologically relevant manner. Two microbial metabolites were administered to illustrate the model’s utility and to highlight the importance of liver first pass metabolism when examining microbial metabolite connections to host physiology and disease. Hepatic first pass metabolism (also known as liver first pass effect) refers to the degree to which a molecule delivered to the liver via the portal vein is transformed prior to entrance into the systemic circulation [Pond and Tozer, 1984]. The higher a molecule’s first pass metabolism, the lower peripheral extra-hepatic tissue exposure. Hence, administration of gut-derived microbial metabolites with high first pass metabolism benefit from portal vein delivery as peripheral (intravenous, subcutaneous, etc.) administration is non-physiologic. Trimethylamine is a well characterized microbial metabolite and of strong interest in our laboratory [Roberts et al., 2018, Schugar et al., 2017, Zhu et al., 2016, Pathak et al., 2020, Schugar and Brown, 2015]. TMA’s conversion to trimethylamine-N-oxide (TMAO) is predominantly catalyzed by hepatic flavin monooxygenase 3 (FMO3), and hence undergoes a degree of liver first pass metabolism. Pharmacokinetic studies investigating TMA metabolism performed in male Wistar rats observed a two-hour half-life and relatively low liver first pass metabolism, demonstrating the majority of TMA within the portal vein is delivered to the systemic circulation [Jaworska et al, 2017; Nnane and Damani, 2001]. As such, portal vein delivery of TMA increases peripheral plasma concentrations, making it well suited for a proof of concept experiment. We observed appreciable rises in both TMA and TMAO peripheral plasma concentrations when administered via portal vein to 8-week C57BL/6 male and female mice (Figure 3). It is also important to note that plasma TMA and TMAO levels were sexually dimorphic, which is in agreement with previous reports [Wang et al., 2011, Bennett et al., 2013]. Compared to non-operated and sham controls, no appreciable differences in dietary intake, weight gain, and motility were observed after the third postoperative day.

The metabolite, 4-hydroxyphenylacetic acid is one of several phenolic acids generated from gut microbial polyphenol and amino acid metabolism [Griffiths and Smith, 1972, Winter et al., 1991]. Preliminary studies demonstrated its relatively high hepatic first pass effect (data not shown), with an appreciable proportion undergoing glucuronidation or sulfation. As such, 4-HPAA would greatly benefit from portal vein over peripheral administration. As expected, 4-HPAA administration did not produce differences in peripheral free or glucuronidated/sulfated fractions of 4-HPAA compared to normal saline vehicle controls (Figure 4C). However, liver levels of total 4-HPAA were increased (Figure 4D). These results highlight the importance of direct intraportal delivery to achieve a physiologically relevant model.

The study of gut microbes in human health and disease is rapidly expanding. Portal vein delivery and hepatic first pass metabolism is a fundamental aspect of many gut microbe-derived metabolites. This method will allow for interrogation of gut microbial metabolites and their links to disease in genetically tractable mouse models. Leveraging this portal vein infusion approach in mice genetically lacking suspected host receptors will be a powerful manner to identify new microbe host signaling pathways relevant to disease pathogenesis [Chkravarthy et al (2009)]. A special related application of this method is the interrogation of bacterial and host lipid signaling given their rapid metabolism when delivered from the gut to liver, making intraperitoneal and other forms of systemic administration less relevant. Other pharmacokinetic and xenobiotic metabolism beyond only microbe-derived metabolite studies will also benefit from applications of this technique. The enterohepatic circulation involves a unique niche of portal vein-enriched metabolites often mediating gut-liver crosstalk to generate vast and largely understudied physiologic effects. This surgical approach will be especially useful in selectively modifying metabolites in this portal blood niche to examine the long-term changes in mouse models of disease.

## Materials and Methods

### Surgical Method

NOTE: Strict adherence to sterile technique is crucial, as contamination may compromise pump and catheter contents.

#### Specialized materials

– Leica Wild M650 surgical microscope; operation performed under 6x – 25x magnification
– Table top rodent anesthesia machine with chamber and nose cone (Parkland Scientific, item no V3000PK)
– Rat intrathecal catheter (short) (Alzet order no. 0007741); distal most flexible portion is 0.36 mm outer diameter (OD), 0.18 mm inner diameter (ID); middle less flexible portion is 0.84 mm OD, 0.36 mm ID; proximal least flexible portion is 1.02 OD, 0.61 mm ID; includes Teflon coated stylet/guidewire; NOTE smaller diameter catheters led to clot formation within the catheter tip lumen
– Polyethylene tubing (Alzet order no 0007750)
– Osmotic pump (Alzet model no 2004); rate of infusion 0.25 uL/hr, up to 4 weeks duration
– Infrared warming pad (Kent Scientific order no DCT-15)
– 10-0 nylon micro suture (AroSurgical item no T04A10N07-13)
– 4.75” × 4” transparent film dressing (3M Tegaderm item no 1626W); small drape with adhesive aperture (3M item no 1092)

#### Filling and priming of osmotic pump

1. A sterile drape is laid within a laminar flow hood.
2. The osmotic pump, polyethylene tubing, and filler needle (provided by Alzet) are placed on the drape, along with a 1 mL syringe.
3. A vial of solution is opened.
4. Sterile gloves are donned. Remove the flow modulator portion of the osmotic pump. Using the syringe and filler needle (ensure no air bubbles are within the needle or syringe), the osmotic pump is slowly filled and the precise amount of the solution documented. Re-insert the flow modulator. Attach the polyethylene tubing (not the catheter) to the pump.
5. Place the pump with attached tubing in a small saline-filled beaker, ensuring the pump is completely immersed. Drape the tubing over the side of the beaker. Cover the beaker with aluminum foil and place in a 37-degree Celsius incubator for 40 hours (specific to pump model no 2004).

#### Surgical method

1. Induction of anesthesia is achieved with 3-4% isoflurane.
2. The abdomen and flank of the mouse are shaved. Sterile eye ointment is applied.
3. The mouse is placed on an infrared warming pad. Anesthesia is maintained with 2-3% isoflurane administered via nose cone. The mouse is secured to the warming pad with silk tape. NOTE the upper extremities should be taped anterior on the nose cone to allow room for later placement of the osmotic pump in the interscapular space.
4. The abdomen is sterilized using 10% betadine and 70% ethanol. Buprenorphine 0.1 mg/kg is administered subcutaneously. Bupivacaine 6 mg/kg is administered subcutaneously along the planned incision.
5. A sterile transparent adhesive film and procedure drape are applied (see materials above). The catheter and instruments are placed on the field.
6. The catheter is prepared by cutting the distal, smallest diameter portion to about **2.5 cm, or one-third the length of the mouse. The guidewire is then cut to be slightly longer than the catheter, and placed within the catheter lumen to protrude about 1 mm from its distal tip.
7. A midline vertical skin incision is made, and the surrounding subcutaneous plane dissected to aid with exposure.
8. The subcutaneous plane of the right lower quadrant is dissected free. A curved hemostat is then used to dissect the subcutaneous plane from the right lower quadrant to the interscapular space on the dorsal side of the mouse; minimal resistance should be encountered. A stitch using 6-0 silk suture is placed within the dermis of the right flank, and its tails left long to be used as a future catheter anchor stitch.
9. A laparotomy is performed from the suprapubic region to the xiphoid process, dissecting the xiphoid free of its lateral attachments.
10. The intestines are externalized leftward and wrapped in moistened gauze (Figure 1B). NOTE prevent twisting of the mesentery to maintain intestinal arterial and venous flow. A stitch using a 6-0 silk suture is placed on the peritoneal/interior-side of the left lower quadrant abdominal wall, and its tails left long to be used as a future catheter anchor stitch.
11. The duodenum is rotated laterally to expose the portal vein (Figure 1B). The left lobe of the liver is externalized superiorly over the xiphoid process and held in place with moistened gauze to expose the liver hilum (Figure 1C). NOTE a moistened gauze and the pressure of a curved hemostat on the duodenum may aid in maintaining exposure.
12. The portal vein is carefully dissected near the splenic vein – superior mesenteric vein confluence. A vessel loop is placed just distal to the confluence using 6-0 silk suture, and left loose. NOTE care should be taken to minimize violation of the pancreas.
13. Approximately 3 mm distal to the vessel loop, a stitch is placed in the anterior one-fourth of the portal vein using a 10-0 nylon micro suture (Figures 1D, 1E). The suture tails and needle are not cut. NOTE the distal portal vein may be gently compressed with forceps to dilate the vein proximally and assist with placement of this stitch.
14. The needle of the prior step is passed through the anterior one-fourth of the portal vein 1 mm proximal to the stitch placed in the previous step (Figure 1F). No knot is tied.
15. The vessel loop is gently pulled inferiorly to occlude flow and produce slight tension on the vein. The catheter-guidewire apparatus is inserted between the stitch and suture placed in steps 13 and 14, respectively (Figure 1G). The catheter is advanced about 5 mm, and the guidewire retracted about 1 cm. NOTE the knot placed in step 13 may be retracted anteriorly slightly to aid in catheter-guidewire insertion. Minimal bleeding should occur around the catheter insertion site.
16. The catheter is pulled back such that its tip protrudes approximately 3 mm within the vein. NOTE further insertion of the cannula increases the risk of clot formation and portal vein occlusion. The free ends of suture in steps 13 and 14 are tied to secure the portal vein around the catheter with such force to slightly crimp the catheter (Figure 1H). The vessel loop is loosened and cut. NOTE portal vein occlusion time should be under 5 minutes. NOTE surgical glue may be used to secure the catheter in place of micro suture, though glue is often non-sterile, produces an inflammatory reaction with dense adhesion formation, and affects gut peristalsis if accidentally applied on bowel.
17. The free tails of the suture from step 16 are wrapped around the catheter twice and tied to create a “Chinese finger trap” effect (Figure 1I).
18. Two stitches are placed to anchor the catheter to the mesentery of the duodenum (Figure 1J). The mesentery should be freed from any retraction, and no tension should be present on the catheter – portal vein junction; otherwise, the position of the mesentery anchor stitches should be adjusted. The catheter should take a path parallel to the natural trajectory of the portal vein, as pressure of the catheter against the sidewall of the portal vein will lead to erosion into the vein sidewall or adjacent tissues. NOTE glue may be used in place of suture to anchor the catheter along the duodenal mesentery, though glue has the aforementioned disadvantages.
19. The guidewire is fully removed. A 22G needle attached to a 1 mL syringe is placed in the lumen of the catheter. Blood is aspirated to remove air from the catheter. About 100 uL of saline is then slowly infused to clear the catheter of blood, and the catheter is clamped at its proximal position.
20. The 22G needle is used to pierce the external left lower quadrant of the abdominal wall (not including the skin) near the anchor stitch placed in step 10. The catheter is then pulled over the needle, and the needle-catheter withdrawn to achieve externalization of the catheter from the left lower quadrant (Figure 1J). NOTE the final orientation of the catheter should be such that no twists are present along its trajectory into the portal vein. The catheter is clamped at its proximal position. The anchor stitch placed in step 10 is then tied around the catheter to secure it to the abdominal wall.
21. The duodenum, intestines, then liver should be placed back into the abdomen in proper anatomic orientation. Final catheter patency and position are checked using the 22G needle to gently aspirate portal blood and ensure smooth infusion of saline. The catheter is again clamped, and attached to the osmotic pump. NOTE slight retrograde portal blood flow into the distal one-third of the catheter is often observed, and will not clot the catheter.
22. Ketoprofen (5 mg/kg) is administered into the peritoneal cavity, along with 0.5 – 1 mL of saline to replace evaporative losses and any blood loss.
23. The abdominal wall is closed with a simple continuous 6-0 polydioxanone (PDS) suture, and the suture tails of the inferior aspect of the closure left long. The catheter is then wrapped across midline, and anchored using the suture tails from the inferior aspect of the abdominal wall closure (Figure 1K).
24. The osmotic pump is placed in the interscapular subcutaneous space using the tunnel created in step 8 (Figures 1K, 1L), and secured with the anchor stitch placed in step 8.
25. The skin is closed in an interrupted horizontal-mattress fashion using a 5-0 polypropylene suture (Figure 1L).

#### Post-operative monitoring and care

1. The mouse is placed on a warming pad to recover, then single-housed to decrease chance of catheter displacement. Water and food are immediately offered, and moistened food pellets are placed on the cage floor.
2. Buprenorphine 0.1 mg/kg is administered subcutaneously on the evening of surgery, and twice daily on postoperative days 1 and 2. Scruffing should be avoided to avoid worsening incisional pain, and displacement of the pump and catheter; rather, subcutaneous injection on the left dorsum of the mouse may be used for medication administration.
3. Twice daily monitoring for three days, and daily monitoring for 14 days are recommended. Special attention should be paid to pain status, activity level, hydration status, incision integrity, pump location, and passage of feces. Signs of distress such as severe pain and markedly decreased activity suggest portal vein occlusion, in which case immediate euthanasia should be performed.
4. Sutures are removed after 14 days.
5. **Necropsy method** (see Supplementary video 6)

### Necropsy method

1. Induction and maintenance of anesthesia is achieved with 4% and 2% isoflurane, respectively. Adequate depth of anesthesia is confirmed by no response to toe-pinch.
2. Isoflurane anesthesia is maintained with a nose cone device, and the mouse is secured to a necropsy board.
3. Laparotomy and thoracotomy are performed.
4. Blood collection via cardiac puncture is performed. NOTE this should be performed prior to testing catheter placement and patency.
5. The catheter is then dissected free and cut transversely near the inferior aspect of the previous abdominal wall closure. A 26G needle attached to a 1 mL syringe of normal saline (devoid of air) is placed within the distal lumen of the freshly cut catheter. Using forceps to compress the catheter tubing around the catheter, saline is gently injected. If the catheter is patent and its tip correctly within the portal vein, the liver parenchyma will blanch.
6. The remainder of the necropsy and tissue collection is at the discretion of the surgeon, though the catheter tip location, portal vein, liver hilum, pancreas, and mesentery of the duodenum should be inspected for any abnormalities.
7. The amount of solution within the osmotic pump should be documented and compared to the amount of solution originally used to fill the pump. Additionally, pump contents may be taken here for analysis to ensure integrity of the solution over the study period.

### Animals

8-10 week male and female C57BL/6J mice were purchased from Jackson Laboratory and housed under specific pathogen–free conditions at the Biological Resources Unit within the Cleveland Clinic Lerner Research Institute, Cleveland, OH. Mice were given standard rodent chow and drinking water ad libitum. Induction and maintenance of anesthesia is achieved with 4% and 2% isoflurane, respectively. Post-surgery, mice were single-housed and given standard rodent chow and drinking water ad libitum. Perioperative mortality was less than 10%. All experiments and procedures were approved by the Institutional Animal Care and Use Committee.

### Reagents

Trimethylamine hydrochloride (Sigma product no T72761) and 4-hydroxyphenylacetic acid (Alfa Aesar product no A15018) were dissolved in 0.9% normal saline (Baxter product no 2F7124) at 9.87 molar and 55 millimolar respectively. These concentrations were based on flow rate and pharmacokinetic calculations aimed to raise plasma concentrations above the normal physiologic level. Before loading into the osmotic pump, each metabolite suspension was sterilized using a 22-micron pore vacuum filtration system.

### Ultrasound

A Vevo 2100 (Visual Sonics) ultrasound was used for ultrasound imaging. Induction and maintenance of anesthesia is achieved with 4% and 2% isoflurane, respectively. After placement of the mouse on a warmed platform, hair removal cream is applied and gently removed with paper towel and liberal application of sterile water. Ultrasound gel is applied, and an MS550S transducer used to image the portal vein and catheter within. Doppler imaging may be used to determine blood flow directionality, as well as turbulence around the catheter. After imaging is complete the ultrasound gel is removed, and the mouse placed in a cage atop a warming pad until awake and recovered.

### Quantification of Trimethylamine (TMA) and Trimethylamine-N-Oxide (TMAO) in Acidified Plasma

Stable isotope dilution high performance liquid chromatography with on-line tandem mass spectrometry (LC–MS/MS) was used for quantification of levels of TMA and TMAO as previously described [Wang et al., 2011, Wang et al., 2014]. Their d9(methyl)-isotopologues, d9-TMA and d9-TMAO, were spiked to plasma, were spiked into plasma as internal standards. LC–MS/MS analyses were performed on a Shimadzu 8050 triple quadrupole mass spectrometer. TMA, TMAO, d9-TMA, and d9-TMAO were monitored using multiple reaction monitoring of precursor and characteristic product ions as follows: m/z 60.2 → 44.2 for TMA; m/z 69.0 → 49.1 for d9-TMA; m/z 76.0 → 58.1 for TMAO; m/z 85.0 → 66.2 for d9-TMAO.

### Sample Preparation and Quantification of 4-hydroxyphenylacetic acid (4-HPAA)

In order to deconjugate plasma samples, 20 uL was treated with β-Glucuronidase/Arylsulfatase obtained from *Helix pomatia* according to the manufacturer’s recommendation (Roche Catalog No. 10127698001). Liver was extracted by homogenizing a piece of liver with 1 mm stainless steel beads at 30 Hz for 5 minutes in methanol + 0.1 % acetic acid. 20 uL of the organic liver extract was dried down under vacuum and resuspended in 20 uL 0.1 M acetate buffer (pH 5.5) and deconjugated according to the manufacturer’s recommendation as outlined above. Stable isotope dilution high performance liquid chromatography with on-line tandem mass spectrometry (LC–MS/MS) was used for quantification of levels of 4-HPAA in plasma, liver, and osmotic pump contents following necropsy. The d6(methyl)-isotopologue of 4-HPAA was used as an internal standard (CDN Isotopes Product No. D-7842). LC–MS/MS analyses were performed using an AB SCIEX Q-Trap 4000 triple quadrupole mass spectrometer equipped with an electrospray ionization source operating in negative ion mode. 4-HPAA was monitored using multiple reaction monitoring of precursor and characteristic product ions as follows: m/z 151.2 → 107.0 for 4-HPAA; m/z 157.2 → 113.0 for d6-4-HPAA. Mass spectrometry parameters were as follows: ions spray voltage −4200 V, ion source heater temperature 400°C, source gas 1 20 psi, source gas 2 30 psi, and curtain gas 20 psi. Nitrogen gas was used for the nebulizer, curtain and collision gas. The HPLC system consisted of four binary pumps (LC-20 AD), autosampler operating at 10°C (Nexera X2 SIL-30AC), and controller (CBM-20A) (Shimadzu Scientific Instruments, Inc.). Chromatographic separations were performed on a reverse phase column (Kinetix XB-C18, 2.6 μm, 150 mm × 4.6 mm ID; Phenomenex, Part No. 00F-4496-E0). Mobile phase A was 5 mM ammonium acetate and 0.1% acetic acid in water and mobile phase B was 0.1% acetic acid in methanol:acetonitrile (9:1; v/v). Samples were injected (10 μl) onto the column equilibrated in 100% A, and separated using a gradient as follows: 0-4 min 0% B, 4-20 min 0-60% B, 20-23min 60-100% B. Flow rate was programmed as follows: 0.3 mL/min. Samples are introduced to the mass spectrometer for analysis from 0-6 min.

### RNA Sequencing in Mouse Tissues

RNA was isolated via the RNeasy Plus Mini Kit (Qiagen) from mouse liver following the manufacturer’s protocol. RNA samples were checked for quality and quantity using the Bio-analyzer (Agilent). RNA-SEQ libraries were generated using the Illumina mRNA TruSEQ Directional library kit and sequenced using an Illumina NovaSeq 6000 (both according to the Manufacturer’s instructions). RNA sequencing was performed by the University of Chicago Genomics Facility. Raw sequence files will be deposited in the Sequence Read Archive before publication (SRA). Paired-end 100 bp reads were controlled for quality with FastQC (v0.11.9, https://www.bioinformatics.babraham.ac.uk/projects/fastqc/) before alignment to the Mus musculus reference transcriptome (NCBI GRCm38.p6 transcript annotations downloaded August, 2020). Reads were aligned using the Kallisto pseudo-alignment (v0.46.0.4, https://pachterlab.github.io/kallisto/) [Bray et al., 2016] with standard parameters using Galaxy (https://galaxyproject.org/) [Afgan et al., 2018]. Pseudo counts were loaded into R (http://www.R-project.org/) (R Development Core Team, 2015) and DESeq2 [Love et al., 2014] (v.1.28.1, https://bioconductor.org/packages/release/bioc/html/DESeq2.html) was used to perform differential expression (DE) analysis on genes with 10 counts per sample with alpha set to 0.05. *P*-values were adjusted using the Benjamini-Hochberg method and genes with *p*<0.05 were considered statistically significant. Heat maps were generated of top 50 differentially expressed transcripts using pheatmap [Kolde 2012] and RColorBrewer [Neuwirth, 2011]. Non-metric multidimensional scaling (NMDS) analysis was performed using the plotMDS function of edgeR [Robinson et al., 2010] using the top 500 DE genes as sorted by log2 fold change. Pathway analysis on the top 250 DE genes was performed using Metascape [Zhou et al., 2019]. The data discussed in this publication have been deposited in NCBI’s Gene Expression Omnibus (Edgar et al., 2002) and are accessible through GEO Series accession number GSE159775 (https://www.ncbi.nlm.nih.gov/geo/query/acc.cgi?acc=GSE159775).

## Acknowledgements

This work was supported in part by National Institutes of Health grants R01 HL120679 (J.M.B.), P50 AA024333 (J.M.B.), and P01 HL147823 (J.M.B.). Use of the Vevo 2100 (Visual Sonics) ultrasound was supported by National Institutes of Health small instrumentation grant 1S10ODO21561.

## Competing Financial Interests

All authors declare no competing financial interests.

## Author Contributions

D.O., L.J.O. and J.M.B. planned the project, designed experiments, analyzed data, and wrote the first draft of the manuscript; F.A., J.C., and Z.W. helped design experiments and provided useful discussion directing collaborative aspects of the project; D.O., L.J.O., K.F., I.C., B.D., Z.W. either conducted mouse experiments, performed biochemical workup of mouse and human tissues, analyzed data, or aided in manuscript preparation; All authors were involved in the editing of the final manuscript.

## Supplementary Material

Supplementary video 1 – surgery part 1

https://drive.google.com/file/d/18sr1wI4dXXoDmTqeVaDleJDrv5rjWyib/view?usp=sharing

Supplementary video 2 – surgery part 2

https://drive.google.com/file/d/1C6Ai9RFhrZb8XZ1bzIKsqaPV_PVSdNm7/view?usp=sharing

Supplementary video 3 – surgery part 3

https://drive.google.com/file/d/1mrKXaKpFufgVt9Lw15YgKHz4usvOAgQl/view?usp=sharing

Supplementary video 4 – ultrasound cannula entry

https://drive.google.com/file/d/1dbKQWan9FEZQhZy8OKUoamPD8ireFnln/view?usp=sharing

Supplementary video 5 – ultrasound cannula doppler

https://drive.google.com/file/d/1GpRsiIWhVD58p5iR_w03tI0w9tgQifkI/view?usp=sharing

Supplementary video 6 – necropsy

https://drive.google.com/file/d/1KgqQIHx4zbAf_BPryx8iZbzq6SHwofKK/view?usp=sharing

## Notes

### Competing Interest Statement

The authors have declared no competing interest.

